# Reintroduction of wild song culture to a critically endangered songbird

**DOI:** 10.1101/2024.11.19.624418

**Authors:** Daniel Lloyd Appleby, Naomi E Langmore, Benjamin Pitcher, Robert Heinsohn, Joy Tripovich, Richard Matkovics, Ross Alexander Crates

**Affiliations:** Fenner School of Environment and Society, Australian National University; Research School of Biology, Australian National University; Macquarie University; University of New South Wales; Taronga Zoo

## Abstract

Animal cultures are learned behaviours, traditions, and collective knowledge that are maintained within populations through social learning. Global biodiversity decline can lead to the loss of animal culture within small and sparsely distributed populations, making the conservation of animal cultures increasingly important. The Critically Endangered regent honeyeater (*Anthochaera phrygia)* is an Australian songbird whose population is declining to the extent that song culture is being lost in the wild. Reintroduced, zoo-bred males that supplement the wild population sing songs that differ from all wild birds, representing a significant cultural barrier that may impact their fitness after release to the wild. Over three breeding seasons within the applied breeding system, we undertook adaptive song tutoring experiments using combinations of song broadcast and live tutoring from two wild-origin males to teach zoo-bred juveniles the wild song. The proportion of juveniles that learned the wild song increased from 0 before the experiment to 42% after three years. The entire population is predicted to sing the wild song within two years. The full version of the wild song taught to zoo-bred males disappeared from the wild over the course of the experiment, making the zoo population the only remaining repository of traditional song culture. Using just two wild founders, we show how animal cultures can be restored in ex-situ populations with simple modifications to husbandry protocols. Ex-situ populations can then play important roles in the maintenance and restoration of wild animal cultures through reintroductions.

## INTRODUCTION

To address the global biodiversity crisis, ex-situ breeding of animals for release into the wild is now an integral conservation tool (Bajomi et al., 2010). The success of animal reintroductions, however, has been the subject of much scrutiny(Fischer & Lindenmayer, 2000; Seddon et al., 2007, Brichieri-Colombi & Moehrenschalager 2016). Whilst practitioners increasingly apply more rigorous approaches to the management of ex-situ populations and the decision-making process around release planning (IUCN/SSC, 2013, Batson et al., 2015, Ogden et al., 2019), the success of ex-situ reintroduction programs in terms of their contribution to recovery of wild populations remains questionable (Bubac et al., 2019).

The suite of factors known to affect reintroduction success is large and still growing (Fischer & Lindenmayer, 2000; Seddon et al., 2007; Berger-Tal et al., 2020). Whilst these factors are often species-specific and linked to whether the initial drivers of population decline have been mitigated (Caughley et 1994, Berger-Tal et al., 2020, Fischer & Lindenmayer 2000, Seddon et al., 2007), there is increasing evidence that the phenotypes and associated post-release fitness of zoo-bred animals can be impacted negatively by life in captivity (Crates et a. 2023). This in turn can hinder the contribution of zoo-bred animals to the recovery of wild populations (Bradley et al., 2014, Lewis, 2021, Appleby et al., 2023).

The predominant focus of ex-situ breeding programs has traditionally been on genetic diversity and disease prevention (Robert, 2009; Williams & Hoffman 2009, Viggers, Lindenmayer & Spratt 1993). Whilst both factors are fundamental for maintaining viable ex-situ populations, behavioural differences between zoo-bred and wild animals rank among the top reported issues hindering the success of reintroduction programs (Berger-Tal et al., 2020). Behaviours play a critical role in the post-release adaptation and survival of ex-situ-bred individuals (Bell, 2016). Issues related to behaviour (Greggor et al., 2016), including social integration (Greggor et al., 2023), foraging skills (Matthews et al., 2005), predator avoidance (McPhee, 2004), dispersal (Berger-Tal 2016) and habitat utilization (Berry et al., 2019) frequently arise during reintroduction programs and can impact the overall success of released populations.

The causes of behavioural differences between ex-situ-bred and wild animals may be in response to either the physical or social environment (Palagi & Bergman, 2021). Manipulating behaviours to improve translocation success is an emerging tool for adaptive management of translocation programs (Greggor et al., 2023; Shier & Swaisgood 2012; Ross et al., 2019). Such manipulations have largely focused on pre-release exposure to novel-stimuli such as antipredator training, or soft-release strategies where individuals can learn and adapt behaviours to novel-environments before being released fully. Culturally acquired behaviours that may impact translocation success have received relatively little focus in translocation planning (Greggor & Goldenberg 2023) but are increasingly being recognised as a critical component of effective conservation (Brakes et al. 2019, Whitten, 2021).

Animal culture refers to the transmission and maintenance of learned behaviours, traditions, and knowledge within animal populations through social learning processes such as observation, imitation, and teaching (Brakes et al., 2019, Brakes et al., 2021). Similar to human culture, animal cultures encompass a wide range of behaviours, including feeding techniques, communication signals, mating rituals, and tool use, which are transmitted within and across generations and may vary between different social groups or populations (Brakes et al., 2019; van de Waal, 2013). These cultural behaviours often contribute to the adaptation and survival of individuals within their respective environments and can shape social dynamics and ecological interactions within animal communities. In captivity, culture may be degraded or lost by preventing necessary social interactions and exposure to suitable adults from which juveniles learn key behaviours (Crates et al. 2023). This is problematic where ex-situ populations contain candidate individuals for release into the wild to reintroduce or wild population supplementation, as culturally-acquired behaviours may be critical for post-release fitness (Brakes et al., 2019). Very few studies have attempted to manipulate culturally learned behaviours in captivity (e.g., Shier and Owlings, 2006; Teitelbaum, Converse & Mueller, 2018).

The regent honeyeater (*Anthochaera phrygia*) is a Critically Endangered Australian songbird (IUCN 2023) with an estimated wild population of fewer than 250 individuals (Garnett & Baker, 2021). To mediate the risk of imminent extinction, zoo breeding to bolster the wild population through reintroductions and provide an insurance against further population decline is a high priority recovery strategy (Commonwealth of Australia, 2016). Supplementation of the wild population with zoo-bred birds has had limited success in arresting the population decline (Heinsohn et al., 2022; Tripovich et al., 2021), with the majority of zoo-bred birds failing to establish a breeding territory or pair with wild mates (Kvistad et al., 2015; Taylor et al., 2018, Tripovich et al., 2021). Historically, this could be explained by the small number of wild birds persisting at the traditional reintroduction site on the southern fringe of the species’ contemporary range in northern Victoria (Kvistad et al., 2015), but the reintroduction strategy shifted in 2021 such that zoo-bred birds are now reintroduced into the greater Blue Mountains in New South Wales (NSW), where most wild regent honeyeaters persist (Heinsohn et al., 2022). Despite this shift, zoo-bred birds still show relatively poor breeding success compared to their wild counterparts in the first breeding season post-release(unpublished data).

Wild male regent honeyeaters sing only one of several song types with loose geographic dialects (Crates et al. 2021). The dominant wild song type within the core range of the remaining wild population is defined as the ‘typical Blue Mountains’ song (Crates et al. 2021). A severe decline in the size and density of the remaining wild regent honeyeater population is leading to loss of the species’ song culture – many young males either learn the songs of other species they encounter or sing an abbreviated version of the ‘typical Blue Mountains’ song with half the number of syllables (Crates et al. 2021, Appleby et al. 2024).

Zoo-bred males sing a distinct abnormal song that is different from all wild song types (Crates et al. 2021). As passerines, young male regent honeyeaters learn their songs via exposure to and imitation of conspecific tutors (Peters, Nowicki & Searcy 2014). The zoo population was founded through the collection of nestlings in 1995 and since then juvenile males have been ‘crèched’ together during their critical song-learning period in early life (Crates et al. 2021). Previously, juvenile birds at Taronga Zoo have been exposed to the songs of wild origin birds through a number playback methods (i.e. Vecsei M, 2015), however these exposures were limited to a single breeding season each (2015, 2019) with all birds in 2015 released into the wild. Zoo-bred regent honeyeaters have therefore never had the opportunity to learn wild regent honeyeater song culture from older conspecifics. Instead, they crystallise an adult song that closely resembles the warbling calls of juveniles, suggesting zoo-bred juveniles learn songs from each other in the creche. The distinct differences between the songs of wild and zoo-bred birds may represent a significant cultural barrier to the assimilation of zoo-bred birds into wild flocks due to assortative mating (Baker, 1983). Indeed, there is evidence of the risk of assortative mating preference within the zoo-bred population; playback experiments show zoo-bred females prefer the familiar but unusual songs of zoo-bred males over the unfamiliar songs of wild males (Appleby et al., 2023). This presents a unique example of an identified wild-ex-situ cultural divide that has direct conservation implications.

We aimed to eliminate an important cultural divide between a zoo-bred and wild animal population within an applied zoo-breeding environment by teaching zoo-bred regent honeyeaters the dominant wild song culture in the area they will be reintroduced to through song tutoring experiments and adaptive management. Our overall goal was to culturally seed the dominant wild song type into the zoo-bred population, such that the wild song culture can easily be sustained in captivity as part of routine management practice.

## Methods

### Study population

The regent honeyeater zoo-breeding program was established in 1995 in response to a dramatic decline in the wild population. 9 nestlings collected from the wild formed the original zoo population, with birds subsequently recruited in 1997 (n=2), 2012 (n = 4), 2019 (n = 3) and 2023 (n = 2). To date, ∼400 zoo-bred regent honeyeaters have been released into the wild.

Birds in this study were housed in Taronga Zoo (TZ), Sydney and/or Taronga Western Plains Zoo (TWPZ), Dubbo. All juveniles were born at either TZ or TWPZ, however, the parents of some juveniles were previously housed at other partner zoos (Adelaide Zoo, Australia Zoo, Australian Reptile Park, Cleland Wildlife Park, Currumbin Wildlife Sanctuary, Featherdale Wildlife Park, Melbourne Zoo, Moonlit Sanctuary Wildlife Conservation Park, Symbio Wildlife Park). All birds included in this study were part of the active zoo-breeding program and therefore subject to routine husbandry protocols. As is often the case in conservation management, resources for time and space within the breeding facilities were limited and as such, experiments needed to be incorporated in the least disruptive manner to ensure they did not negatively impact the breeding capacity of the program.

### Tutoring Protocols

During the experiments birds were housed in one of three aviary types. The first, aviary type 1 at Taronga Zoo, (6m x 3.3m x 3m), the second two at Taronga Western Plains Zoo, aviary type 2 (4m x 4m x 6m) and type 3 (6m x 4m x 6m). Aviaries were typical of those used as crèche aviaries during normal breeding and post-breeding procedures (Fig. 1).

**Figure 1:**
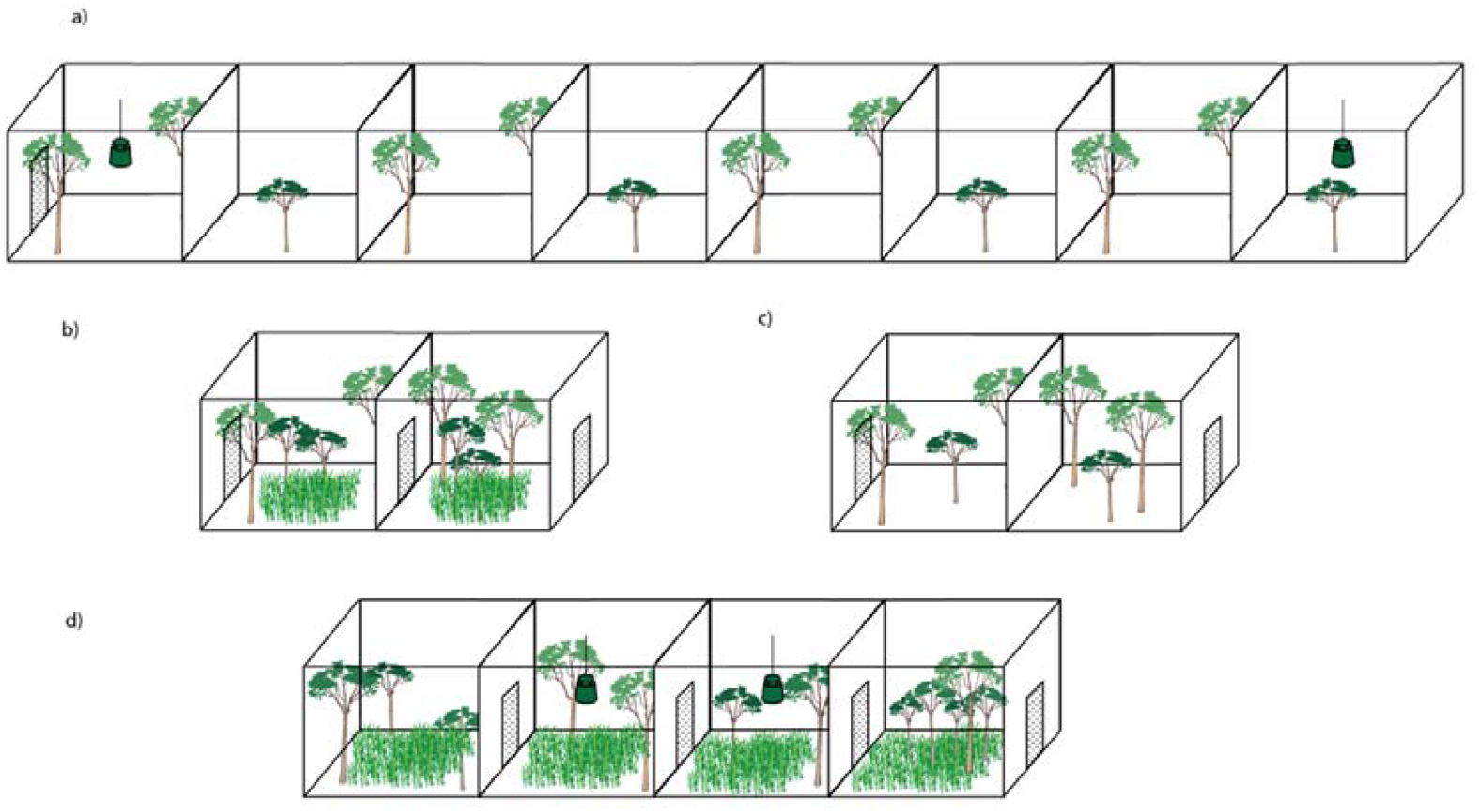
Schematic of regent honeyeater song tutoring aviaries for the different experimental protocols. A) shows playback only tutoring aviary configuration, where juvenile pupil groups were placed in outer aviaries of an eight-aviary block with broadcast speakers. Breeding pairs housed in the aviaries between. Aviaries were separated by solid dividers preventing visual contact between aviaries. B) shows the configuration of the live tutor only cohort (large) where adult tutor was initially housed in the left aviary while juvenile pupils were housed in the right aviary but could both see and hear the tutor. The tutor was subsequently introduced to the juveniles at the conclusion of the breeding period by opening the adjoining door. C) shows the configuration of the live tutor only (small) cohorts where both the adult tutor and juvenile pupils were housed together in one aviary. Two cohorts were housed side by side with a solid divider between preventing visual contact. D) shows the live tutor + playback configuration where the adult tutor was initially housed in the left aviary while juvenile pupils were housed in the right aviary, but could both see and hear the tutor as well as exposed to broadcast playback. The tutor was subsequently introduced to the juveniles at the conclusion of the breeding period by opening the adjoining door.

We conducted tutoring experiments over three Austral post-breeding seasons (September - March) commencing in 2020. We implemented five treatments and one control group across the two breeding institutions in the first season and adaptively refined experiments before each subsequent season based on the results of previous seasons’ experiments.

We considered two common experimental song tutoring techniques; live tutors and audio playback as options for teaching juvenile birds to sing the ‘typical Blue Mountains’ wild song. Live tutoring has shown more success than audio-only tutoring (Catchpole & Slater, 2008; Soma, 2011), however due to the rarity of this species in the wild, acquiring wild birds is a logistical and ethical challenge that is compounded by regional song differences and the presence of wild birds with abnormal songs (Crates et al., 2021). Only two wild males recruited to the TZ breeding population in 2019 sang the ‘typical Blue-Mountains’ song. Unlike some species, (e.g. European starlings *Sturnus vulgaris*, Mountjoy & Lemon, 1994; canary *Serinus canaria*, Nottebohm, 1986), evidence suggests regent honeyeaters are ‘close-ended’ song learners, with males’ songs becoming fixed after a period of learning and crystallisation in the first year of life (Crates et al. 2021). Only these two wild-caught males were available as live tutors in the first year of the experiment.

Playback tutoring, although generally considered inferior to live tutoring, has proven successful in many species (e.g. chaffinches *Fringilla coelebs*; Thorpe, 1958) and possesses several logistical benefits in the zoo-breeding setting. In regent honeyeaters, wild song recordings were readily available from previous research (Crates et al., 2021) and could be implemented inexpensively. Secondly, playback could be implemented with minimal disruption to the zoo’s routine husbandry activities. If successful, playback tutoring could easily be expanded to other facilities involved in the ex-situ breeding program. The remaining years of the study were adapted based on preliminary results of the previous years, the rationale for which can be found in the supplementary text (Text S1).

### Standard timeline

All treatments shared the same timeline, with young males transferred from their natal aviaries to their respective experimental crèche aviaries (Fig. 2) approximately two to three weeks post-fledging (∼30 days old). Juveniles are typically moved out of their natal aviaries at this point as fathers become territorial and aggressive towards juveniles prior to re-nesting (Regent honeyeater husbandry guidelines, Taronga Zoo, 2013). Pairs can have up to three clutches per breeding season (Hibbard, 2011). Juvenile males were exposed to song tutoring from the time they entered the experimental crèche until the conclusion of the breeding season when they are transferred to a larger, multi-species flight aviary, which has been shown to benefit post-release survival (Tripovich et al., 2021). Details of experimental timelines can be found in the supplementary material (Table S1). Some juveniles from second or third broods received a shorter duration of song tutoring (Range = 128 – 208 days, mean = 162), but all were moved into the experimental aviaries at approximately the same age at around 30 days old. We considered this to be reflective of the typical wild song-learning environment where juveniles from different broods join larger, mixed-age post-breeding flocks towards the end of the breeding season (Geering & French 1998). The five treatments and the control were conducted under the following protocols, summarised in Table 1:

**Table 1:**
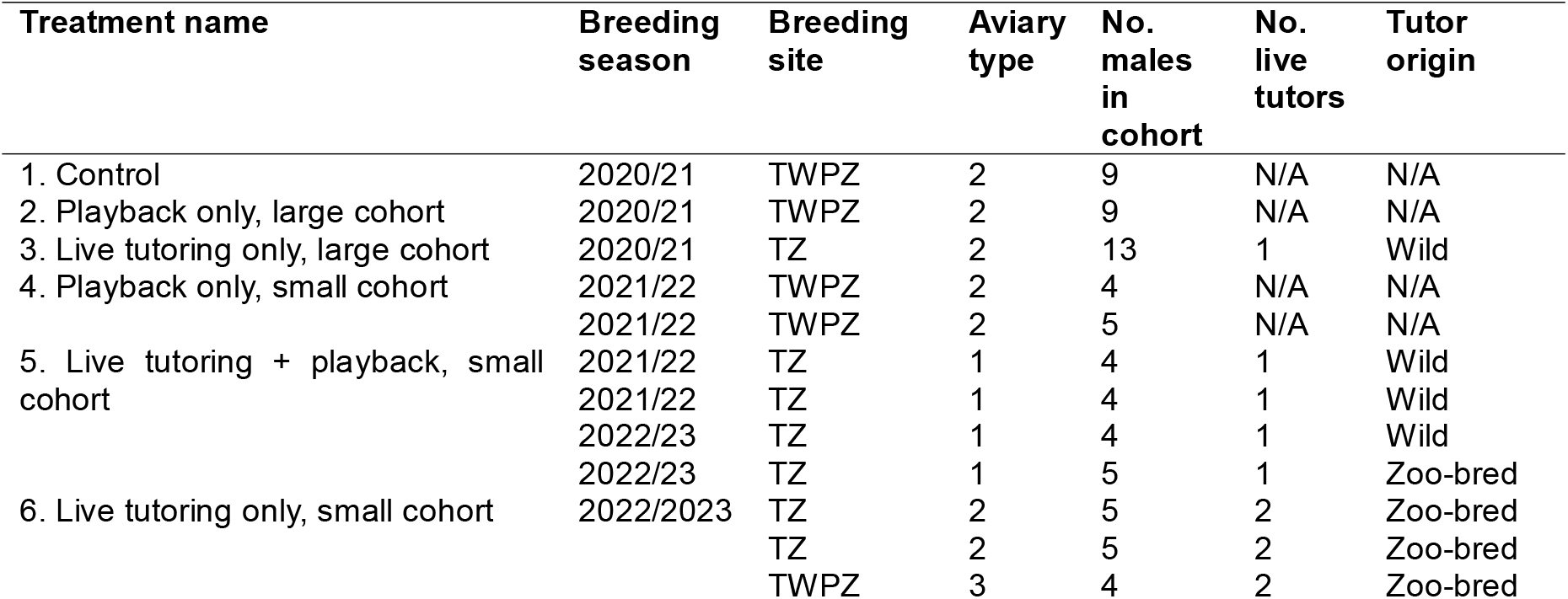
Summary of the zoo-bred regent honeyeater song tutoring experimental design under an adaptive management framework showing cohorts of juveniles, their tutoring treatment, site, aviary type and cohort size. Breeding site TWPZ refers to Taronga Western Plains Zoo, Dubbo. TZ refers to Taronga Zoo, Sydney. Zoo-bred birds in the Tutor origin column refer to zoo-bred males that successfully learned the typical Blue Mountains song in previous years of the study. See Fig. 1 for description of aviary types

**Fig 2:**
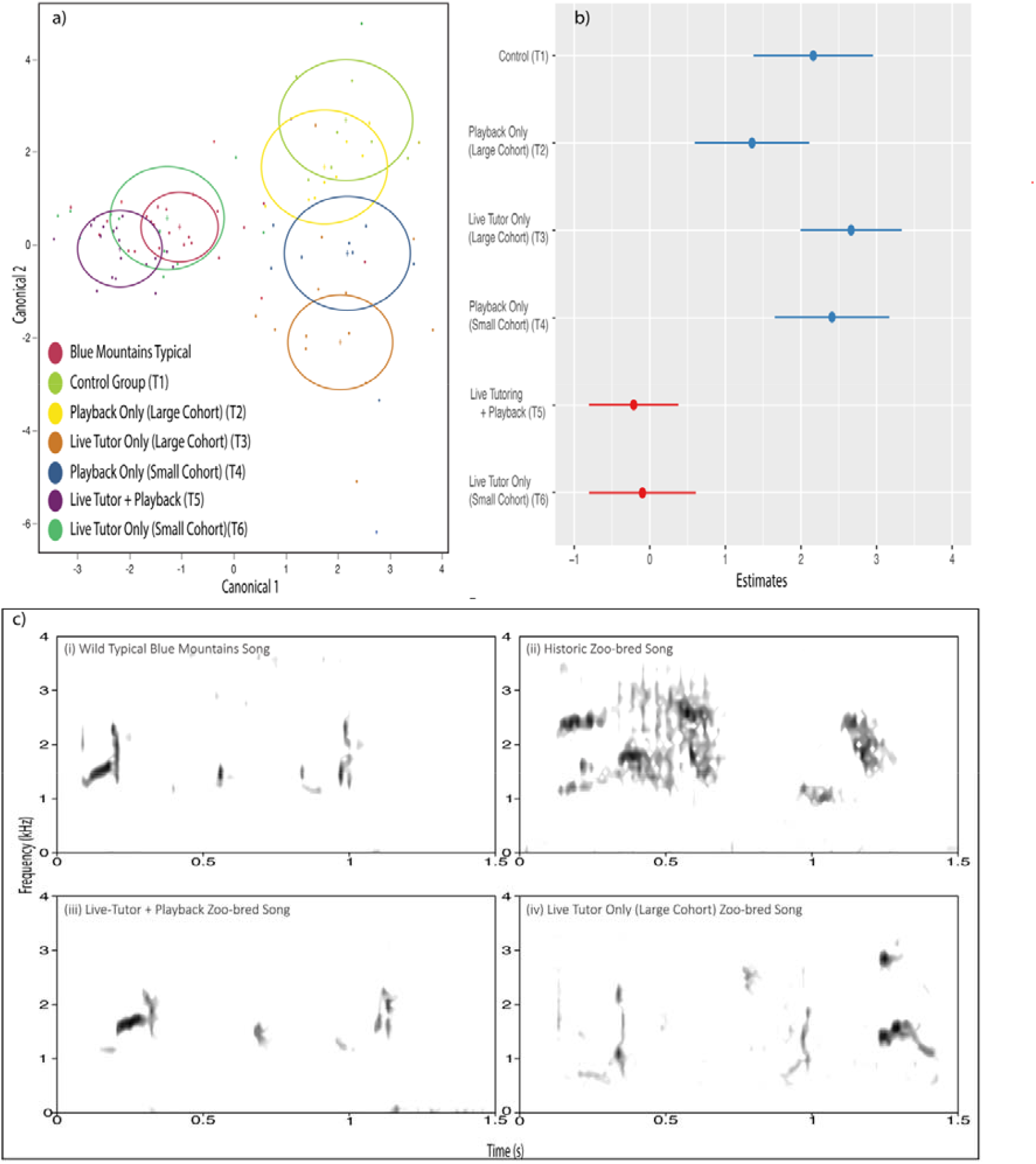
(a) Discriminant function analysis (DFA) of regent honeyeater songs by treatment group. DFA labels each treatment group or reference group multivariate mean with a circle corresponding to a 95% confidence limit for the mean. Groups whose songs are significantly different have nonintersecting circles. See Figure S3 for DFA including the historic captive songs. Model estimates showing the effect of regent honeyeater song tutoring experimental treatment on the similarity of songs to the culturally dominant ‘typical Blue Mountains’ wild regent honeyeater songs based on Mahalanobis distances. Points show predictions, lines show 95% confidence intervals. Estimates in red show the two treatment groups (5 & 6) whose songs show no statistically significant difference to the typical Blue Mountains reference songs as per Table 2.Spectrograms showing example songs of: (i) wild Typical Blue Mountains; (ii) Historic zoo-bred song; (iii) tutored zoo-bred male with live tutor and playback (Treatment 5); (iv) tutored zoo-bred male with live tutor only (large cohort-Treatment 3).

### Treatment 1 - Control

The control tutoring was conducted at TWPZ in the 2020-2021 breeding season. Nine juvenile males were crèched together according to traditional husbandry protocols on the standard timeline described above and in Table S1. Birds were housed in aviary type 2 (Fig. 1a) at the end of a breeding block of aviaries and were unable to see birds in other treatments. The control cohort was separated from the closest treatment cohort (Treatment 2) by 8 aviaries, a total distance exceeding 32 m. A dummy speaker (Bose FreeSpace 360p Series II) was mounted within the aviary but did not emit sound. The only sounds juvenile males in the control group were exposed to were from each other, and from the surrounding environment.

**Table 2:** Model Summary derived from a general linear model showing the effect sizes of treatment group in explaining the differences between the songs of zoo-bred regent honeyeaters. Models are based on the Mahalanobis distance between the mean acoustic attributes of males in each group relative to the average culturally-dominant typical Blue Mountains wild song. P-values in bold indicate significant differences at the p < .05 level.

| *Coefficient* | *Estimate* | *SE* | *t* | *p* |
| --- | --- | --- | --- | --- |
| *Control (T1)* | 2.16 | 0.40 | 11.39 | **<0.001** |
| *Playback Only (Large, T2)* | 1.35 | 0.30 | -0.72 | **<0.001** |
| *Live Tutor (Large, T3)* | 2.66 | 0.34 | 7.91 | **<0.001** |
| *Playback Only (Small, T4)* | 2.41 | 0.38 | 6.33 | **<0.001** |
| *Live Tutoring & Playback (T5)* | -0.21 | 0.30 | -0.72 | 0.48 |
| *Live Tutoring (Small, T6)* | -0.10 | 0.36 | -0.27 | 0.79 |

### Treatment 2 – Playback only (large cohort)

The playback only (large cohort) treatment was conducted at TWPZ in the 2020-2021 breeding season. Nine birds were housed together in an aviary type 2 (Fig. 1a) according to the standard timeline. A composite track consisting of the songs of 25 different wild male regent honeyeaters singing the typical Blue Mountains song type (each with a duration of ∼2 seconds) was broadcast to the juvenile birds from sunrise to sunset every day of the experiment from a broadcast speaker mounted within the aviary. To approximate the natural variation in songbird singing activity (Aide et al., 2013; Bruni et al., 2014), songs were emitted at a more frequent rate (∼four songs per minute) for the first four hours and last two hours of the day, than during the middle of the day (∼one song per minute). The length of the playback track was increased two months into the experiment to reflect the lengthening of the daylight hours during the Austral summer. A description of the process of composing the playback track, playback system parameters and playback track itself can be found in the supplemental text (Text S2 & Text S3).

### Treatment 3 – Live tutoring only (large cohort)

The live tutoring only (large cohort) was conducted at TZ in the 2020-2021 breeding season. Thirteen juvenile males were housed together in an aviary type 1 (Fig. 1b) according to the standard timeline. Juveniles were moved into an aviary adjacent to one of the two wild-origin adult males that sang the typical Blue Mountains wild song. Birds could both see and hear the adult male but could not physically interact with him for the first 2.5 months, during which period the adult male was breeding and would otherwise have shown aggression to juveniles in the same aviary (Regent honeyeater husbandry guidelines, Taronga Zoo, 2013). After 2.5 months, the two aviaries were merged, and the juveniles could directly interact with the adult male.

### Treatment 4 – Playback only (small cohort)

The playback only (small cohort) was conducted at TWPZ in the 2021-2022 breeding season. Four juvenile males were housed in an aviary type 2 (Fig. 1a) according to the standard timeline and were unable to see birds in adjacent aviaries. All other experimental parameters were as described above for Treatment 2-Playback only (large cohort).

### Treatment 5: Live tutoring + playback

The live tutoring + playback treatment was conducted at TZ in the 2021-22 and 2022-2023 breeding seasons. 18 juvenile males were housed in two separate aviaries: eight (2021-2022) in a type 1 aviary (Fig. 1d) and 10 (2022-2023) also in a type 1 aviary (Fig. 1d). Birds were moved from their natal aviary into a crèche aviary adjacent to a wild-origin adult male according to the standard timeline. As in treatment 3, juvenile birds were able to both see and hear the adult male but could not physically interact with him for the first 2.5 months until the aviaries were merged. Unlike treatment 3, during the period that juveniles could not interact with the wild-origin adult male, they were exposed to playback in the same way as treatment 4. After the juveniles and adult male were united, the broadcast playback continued until the end of the breeding season.

### Treatment 6: Live tutoring only (small cohort)

The live tutoring only (small cohort) experiment was conducted at both TZ and TWPZ in the 2022-2023 breeding season. Fourteen birds were housed in three separate aviaries, with five juveniles each in two type 2 aviaries (Fig. 1c) at TZ and four juvenile males in one type 3 aviary at TWPZ (Fig. 1c). In the live tutoring only (small cohort) experiments, juvenile males were transferred from their natal aviary according to the standard timeline to a crèche aviary with a zoo-bred adult male tutor (∼18 months old) that had successfully learned the typical Blue Mountains song in the previous season’s Treatment 5 experiment. During the tutoring period they could see, hear and physically interact with the live tutor.

In summary, the experiment involved 68 birds: 2 wild origin males (tutors) and 66 juvenile zoo-bred males, of which four were used as live adult tutors in subsequent years (Table 1).

### Song recording and editing

Juveniles’ songs were recorded in their tutoring aviaries at the end of each tutoring period (October-March; Age range: 2.5 months – 7 months) before they were moved into a multi-species flight aviary in preparation for potential release to the wild. Songs of each juvenile male were recorded over two days at a typical distance of 1 – 4m using a Sennheiser ME66 microphone and a Zoom H4n Pro recording device (48KHz sampling rate, 16 bit). For comparison with the songs of zoo-bred juveniles included in the tutoring experiments, we also obtained recordings of (i) ‘typical Blue Mountains’ wild birds that were made in the wild between 2015 and 2019 (which formed the playback tracks used for audio tutoring); and (ii) zoo-bred birds that had not been exposed to song tutoring either one week after release into the wild in 2017 (n = 12) or within captivity in 2019 (n = 9). These additional recordings used the methods detailed in Crates et al. (2021). We first edited individual songs (342 total, min 4 max 13 per individual) in Apple Garageband (v10.3.3) to trim them to include only the target individual’s song and no other background noise.

### Spectral and statistical analysis

To assess the similarity between the songs of tutored juveniles and the reference wild-type typical Blue Mountains males, we first performed semi-autonomous syllable segmentation using the open-source Python software ‘Chipper v1.0’ (Searfoss et al., 2020). Chipper allows users to set default parameters for automated segmentation of syllables within each bout of song. We used only the top two percent on signal and used the default settings of 10 ms minimum silence duration and 30 ms minimum syllable duration. Occasionally we adjusted the syllable onsets and offsets (syllable selections) to remove environmental noises or other artefacts from recordings. We subsequently performed Chipper’s song analysis to obtain spectral, song and syllable measurements, from which we selected nine acoustic features of the songs for subsequent analysis (Table S2).

After ensuring none of the attributes showed strong positive or negative correlation with any others using the ‘ggcorrplot’ function in the R-package ‘GGally’ (Schloerke, Crowley & Cook, 2018; Figure S2), we averaged each spectral measure across multiple recordings of each individual’s song. We then performed discriminant function analysis (DFA) on the average spectral measures using JMP v17.2.0 (SAS Institute Inc., Cary, NC). The DFA contained eight pre-defined groups: the five experimental groups and the control group described above, plus the songs of untutored zoo-bred males from previous years and the typical Blue Mountains wild male reference songs, which included songs of the two wild-origin live tutors.

We then calculated the Mahalanobis distance between each individual’s average song attributes and the centroid of the culturally-dominant typical Blue Mountains reference songs recorded in the wild between 2015 and 2017 and used in the playback tutoring experiments. The Mahalanobis distance represents a measure of acoustic (dis)similarity between the average song of each individual regent honeyeater and the target song of the tutoring experiment (De Maesschalcket al., 2000). Increasing Mahalanobis distances indicate increasing dissimilarity between an individual’s learned song and the mean of the culturally dominant wild song type.

To assess whether the songs of birds in each treatment group differed from the typical Blue Mountains reference songs (and from those in other treatment groups), we used the Mahalanobis distances described above as the response variable in a generalised linear model (GLM) using the package ‘stats’ in R 4.0.3 (R core team, 2020). The model contained treatment group as a factorial fixed effect, individual age in days at song recording as a continuous fixed effect. Age was not a significant predictor in this model. We conducted backwards model selection using the ‘stepAIC’ function from the package ‘MASS’ (Venables & Ripley, 2002), which revealed that removing age as a predictor improved our model (Table S3). We selected a generalised linear model containing only treatment as a factorial fixed effect. We could not include breeding facility or aviary type as a random term due to unavoidable restrictions on the space available within the breeding facilities for experimentation. We assessed the quality of the model by using the ‘simulateResiduals’ function in the ‘DHARMa’ package (Hartig, 2022, Fig. S2) and conducted post-hoc tests of pairwise significance between treatment groups using the package ‘multcomp’ v1.14-17 (Hothorn et al., 2016) in R.

## Results

The first two discriminant functions explained over 84% of the total variance in the song samples. The model’s overall significance was supported by multiple test statistics (*wilks-lambda=* 0.062, f = 20.10, p = <0.001; Pillai’s Trace = 1.83, f= 13.90 p= <0.001).

Before the first year of the experiment, only the two wild-origin males sang the typical Blue Mountains song within the zoo population. No juveniles successfully learned the typical Blue Mountains song during the first year of the experiment (Treatments 1 – 3: control group, playback only large group and tutor only large group, Figure 2a). Model predictions based on Mahalanobis distances showed the average songs of males in all three groups were significantly different from the reference group (Figure 2b, Table 2).

By the end of the second season, no juveniles in Treatment 4 (playback only, small group) produced the typical Blue Mountains song, however eight juvenile males in Treatment 5 (live tutor + playback) learned to sing songs that closely-resembled the typical Blue Mountains song type. Discriminant function analysis (DFA) was unable to distinguish the songs of these birds from the wild reference songs (Figure 2a), with the GLM confirming the songs of birds in this group were not significantly different from the reference songs (Figure 2b, Table 2).

Four males from Treatment group 5 were employed as live tutors in the third year of the experiment. Together with the two wild origin birds, these four males successfully tutored an additional 19 juveniles in year three; all within Treatment 6 (live tutor only, small cohort). The songs of these birds could also not be distinguished from the typical Blue Mountains reference songs by DFA (Figure 2a). Again, there were no significant differences between the songs of birds in Treatment 6 and those of the wild reference group (Figure 2b-c). Songs of younger juvenile males hatched later in the season were no less similar to the reference songs than those of older males from first clutches (Table S4).

By the end of the third year of the experiment, 32 males representing approximately ∼42 % of the male zoo population (which fluctuates annually depending on the number bred and the number reintroduced to the wild) sang songs that were statistically indistinguishable from the typical Blue Mountains song based on DFA (Figure 2a). With the observed rate of cultural seeding of the typical Blue Mountains song within the zoo population, we predict all males within the zoo-bred regent honeyeater population will learn to sing the wild song within 3 years.

## Discussion

The importance of considering animal cultures in conservation programs is increasingly being acknowledged (Brakes et al. 2019). We show that it is possible to reintroduce lost wild culture into an ex-situ animal population by providing the first example of successful song tutoring in an active conservation program. Both live tutoring (small cohort) and live tutoring with playback supplementation were successful in training birds to sing songs that did not differ significantly from the reference wild song. We were able to utilise birds from previous year’s experiments to increase the rate at which the wild song culture was culturally seeded in the zoo population. Playback tutoring in both large and small tutor groups was unsuccessful and did not differ significantly from the control. Live tutoring (large cohort) alone did differ from the control but was also significantly different from the reference wild song. These results suggest that being able to interact with a live tutor in a social group with a low ratio of adult tutors to juveniles are the key factors influencing whether birds are able to successfully acquire the wild song culture.

In numerous bird species, tutoring via playback is observed to be less effective than instruction from a live tutor (Soma, 2011; Varkevisser et al., 2022). The understanding of avian learning processes predominantly stems from a limited selection of taxonomic groups. In this study, birds subjected solely to playback tutoring exhibited an expansion in song complexity relative to untutored, zoo-bred birds but failed to replicate accurately the target wild-type song. It has been suggested that birds exclusively exposed to playback tutoring may exhibit lower success rates in learning the reference song (Mets & Brainard, 2018; Varkevisser et al. 2022). As the birds progress into the sensorimotor stage of learning, the songs of associates may provide a more enriched form of learning than the playback (Mets & Brainard, 2018), which could facilitate horizontal transmission of song among juveniles within the aviary. This horizonal transmission would then be compounded by the addition of younger birds throughout the experiment who may learn predominantly from older juveniles rather than the audio playback. This hypothesis aligns with the findings of Deregnaucourt and Gahr (2013), who show that within generation song learning can occur in zebra finches *Taeniopygia castanotis* where a young bird exposed to songs of a tutor can teach another bird not exposed to the tutor some elements of reference song. In that study, juvenile songs converged more closely with each other than with the initial reference tutor.

The live-tutoring only (large cohort) also failed to facilitate juveniles to accurately produce wild-type regent honeyeater songs, although the songs of juvenile males within this group were closer to the wild reference songs than those of the playback treatments. We propose a similar mechanism for this result; in this instance juvenile birds did not receive playback during the earliest stages of independence, but rather had no tutoring until the adult tutor was moved into the experimental aviary. While the exact timing of song learning in this species is unknown, we propose that the introduction of the live tutor fell between the sensory and sensorimotor stages and as such limited the amount of sensory learning opportunity for the juvenile males in Treatment 3. Experiments depriving young birds of tutors have demonstrated that the absence of a tutor in critical learning phases reduces the likelihood that young birds will learn new syllables after the introduction of a tutor (Mackevicius et al., 2023). It is also possible that as the oldest birds were deprived of a live tutor for 2.5 months, their maladapted songs may have then been horizontally transmitted to other birds in the cohort.

The two successful treatments shared two key features: opportunities for multimodal learning from live tutors and small cohorts. There is probably a relationship between smaller cohorts and truly multimodal learning. Where the live tutoring only (large cohort) presented the *opportunity* for multimodal learning, it was probably not realised due to the large ratio of students to tutors. The small cohort size may have allowed for frequent and prolonged interaction with the tutor, and this is consistent with previous research that links the number of tutors with song learning fidelity (Williams & Slater, 1990). Anecdotally, we observed this occurring in real time – in the large cohort group the tutor was overwhelmed by practicing juveniles and frequent song practicing was seen between juveniles (Figure 3). In the small cohorts, the adult male dominated high value areas like the feeders more easily. During these interactions head-bobbing and bill-snapping typically associated with the crescendo of the typical Blue Mountains song (supplemental video 1).

While both successful cohorts were not statistically distinguishable from the reference wild song, birds that only received live tutoring resembled their specific tutor more closely than the wild reference. This is perhaps due to the increased variation in reference songs available to these birds but critically, unlike the playback only cohort, the live tutoring + playback cohort had exposure to a wider range of songs, that may impact the reference template and birds own song experience consistent with the eavesdropping hypothesis (Burt et al., 2007).

While it is possible that natural cultural drift caused the divergence over time between wild and zoo-bred regent honeyeaters, it is highly likely that hand rearing of founder juveniles caused significant cultural divergence and drift within the zoo population (Crates et al., 2021; Appleby et al., 2023). Our results show that although the control group remained the least similar of all treatments to the typical Blue Mountains song, those birds’ songs were considerably more similar to typical wild song than historical ex-situ songs. This may be reflective of the increased socialisation that a larger ex-situ population affords, or may be due to changes to housing arrangements that crèche juvenile birds closer to adult, albeit ex-situ tutors.

The results of our study may have significant implications for regent honeyeater conservation efforts. Regent honeyeaters bred in captivity have generally shown a low propensity to integrate with wild birds after being released and an even lower propensity to pair and breed(Taylor et al., 2018). Assortative mating based on differences in song culture is common amongst birds (Slabbekoorn & Smith, 2002; Price 2008) and previous research suggests that zoo-bred female regent honeyeaters prefer the familiar songs of zoo-bred males to the unfamiliar typical Blue Mountains song produced by most wild males (Appleby et al., 2023). Reintroducing wild songs to the ex-situ population may increase the likelihood of breeding events between reintroduced and wild regent honeyeaters (Appleby et al., 2023) and increase social cohesion at the flock level (Lewis et al., 2021; Martins et al., 2018).

At the inception of this project the typical blue mountains song was the most common song sang in the target release site. Since 2020, monitoring has shown that the typical Blue Mountains song has effectively disappeared from the wild population and has been replaced by the more simplified ‘clipped Blue Mountains’ song type that contains half the number of syllables (Appleby et al. 2024). Whilst cultural drift and revolution are common features of animal populations, it is more likely that simplification of the song culture is reflective of an ongoing decline in the size and density of the wild regent honeyeater population (Heinsohn et al. 2022), making it increasingly challenging for cultural diversity to be maintained at the population level (Crates et al. 2021; Appleby et al. 2024). This decline in song complexity may have downstream Allee effects that negatively impact the viability of the wild population (Paxton et al., 2019; Laiolo et al., 2008). The results of our study mean that zoo-bred birds are now the sole source of traditional song cultural knowledge in regent honeyeaters. As it has been shown previously that songbirds can learn songs experimentally in the wild (Mennill et al., 2018), the results of our study opens up the exciting opportunity to use the reintroduction of zoo-bred birds to re-establish or reinforce the species’ traditional song culture in the wild population.

Our study highlights the important role of experienced adults as repositories of cultural knowledge in conservation programs. It also demonstrates the importance of facilitating social interactions at all stages of the husbandry process in ex-situ breeding, to help maintain socially acquired behaviours that may have a significant bearing on post-release fitness and the overall effectiveness of reintroduction programs (Crates et al. 2023). Given the increasing recognition of the need to consider animal cultures in conservation (Brakes et al. 2021), we provide clear evidence that this can be achieved rapidly and at little cost within applied conservation settings through experimentation and adaptive management. There is great scope for similar studies on other endangered species to help reduce to the phenotypic divide between wild and zoo-bred populations (Crates et al. 2023). This will require improved monitoring of wild animal populations to better understand the dynamics of culturally-acquired behaviours (Scheele et al. 2021), as well as increased collaboration between academics and applied conservation practitioners to facilitate research, reporting and implement evidence-based changes to zoo-husbandry and reintroduction strategy (Crates et al. 2023). Such actions could make a significant contribution to minimising the loss of not only global biodiversity over coming decades, but also its emergent properties such as birdsong that form an intrinsic part of human culture and wellbeing.

## References

Appleby, D. L., Tripovich, J. S., Langmore, N. E., Heinsohn, R., Pitcher, B. J., & Crates, R. (2023). Zoo-bred female birds prefer songs of zoo-bred males: Implications for adaptive management of reintroduction programs. Biological Conservation, 284, 110171. 10.1016/j.biocon.2023.110171

Appleby, D.L., Langmore N.E., Heinsohn R., Crates, R. (2024) Frequency-dependant shifts in song-preference may decrease fitness costs associated with reduced bird song complexity. Royal Society Open Science

Bajomi, B., Pullin, A. S., Stewart, G. B., & Takács-Sánta, A. (2010). Bias and dispersal in the animal reintroduction literature. Oryx, 44(3), 358–365. 10.1017/s0030605310000281

Baker, M. C. (1983). The behavioral response of female Nuttall’s white-crowned sparrows to male song of natal and alien dialects. Behavioral Ecology and Sociobiology, 12, 309–315.

Bates, D., Maechler, M., Bolker, B., & Walker, S. (2015). Fitting Linear Mixed-Effects Models Using lme4. Journal of Statistical Software, 67(1), 1–48. 10.18637/jss.v067.i01

Batson, W. G., Gordon, I. J., Fletcher, D. B., & Manning, A. D. (2015). REVIEW: Translocation tactics: a framework to support the IUCN Guidelines for wildlife translocations and improve the quality of applied methods. Journal of Applied Ecology, 52(6), 1598–1607. 10.1111/1365-2664.12498

Bell, B. D. (2016). Behavior based management: conservation translocations. Conservation behavior: applying behavioral ecology to wildlife conservation and management, 21, 212.

Berger-Tal, O., Blumstein, D. T., Carroll, S., Fisher, R. N., Mesnick, S. L., Owen, M. A., Saltz, D., St. Claire, C. C., & Swaisgood, R. R. (2016). A systematic survey of the integration of animal behavior into conservation. Conservation Biology, 30(4), 744–753. 10.1111/cobi.12654

BergerℒTal, O., Blumstein, D. T., & Swaisgood, R. R. (2020). Conservation translocations: a review of common difficulties and promising directions. Animal Conservation, 23(2), 121–131. 10.1111/acv.12534

Berry, L. E., L’ Hotellier, F. A., Carter, A., Kemp, L., Kavanagh, R. P., & Roshier, D. A. (2019). Patterns of habitat use by three threatened mammals 10ℒyears after reintroduction into a fenced reserve free of introduced predators. Biological Conservation, 230, 1–9. 10.1016/j.biocon.2018.11.023

Bradley, H. S., Tomlinson, S., Craig, M. D., Cross, A. T., & Bateman, P. W. (2022). Mitigation translocation as a management tool. Conservation Biology, 36(1), e13667. 10.1111/cobi.13667

Brakes, P., Dall, S. R. X., Aplin, L. M., Bearhop, S., Carroll, E. L., Ciucci, P., Fishlock, V., Ford, J. K. B., Garland, E. C., Keith, S. A., McGregor, P. K., Mesnick, S. L., Noad, M. J., Notarbartolo di Sciara, G., Robbins, M. M., Simmonds, M. P., Spina, F., Thornton, A., Wade, P. R., Rutz, C. (2019). Animal cultures matter for conservation. Science, 363(6431), 1032–1034. 10.1126/science.aaw3557

BrichieriℒColombi, T. A., & Moehrenschlager, A. (2016). Alignment of threat, effort, and perceived success in North American conservation translocations. Conservation Biology, 30(6), 1159–1172. 10.1111/cobi.12743

Bubac, C. M., Johnson, A. C., Fox, J. A., & Cullingham, C. I. (2019). Conservation translocations and post-release monitoring: Identifying trends in failures, biases, and challenges from around the world. Biological Conservation, 238, 108239. 10.1016/j.biocon.2019.108239

Burt, J. M., O’Loghlen, A. L., Templeton, C. N., Campbell, S. E., & Beecher, M. D. (2007). Assessing the importance of social factors in bird song learning: a test using computerℒsimulated tutors. Ethology, 113(10), 917–925.

Catchpole, C. K. S. P. J. B. (2008). Bird song: Biological themes and variations, second edition. Cambridge University Press. 10.1017/CBO9780511754791

Caughley, G. (1994). Directions in Conservation Biology. Journal of Animal Ecology, 63(2), 215–244. 10.2307/5542

Commonwealth of Australia. (2016). National Recovery Plan for the Regent Honeyeater (Anthochaera phrygia).

Crates, R., Langmore, N., Ranjard, L., Stojanovic, D., Rayner, L., Ingwersen, D., & Heinsohn, R. (2021). Loss of vocal culture and fitness costs in a critically endangered songbird. Proceedings of the Royal Society B: Biological Sciences, 288(1947), 20210225–20210225. 10.1098/rspb.2021.0225

Crates, R., Stojanovic, D., & Heinsohn, R. (2023). The phenotypic costs of captivity. Biological Reviews, 98(2), 434–449. 10.1111/brv.12913

De Maesschalck, R., Jouan-Rimbaud, D., & Massart, D. L. (2000). The mahalanobis distance. Chemometrics and intelligent laboratory systems, 50(1), 1–18.

Derégnaucourt, S., & Gahr, M. (2013). Horizontal transmission of the father’s song in the zebra finch (Taeniopygia guttata). Biology Letters, 9(4), 20130247.

Elsbeth McPhee, M. (2004). Generations in captivity increases behavioral variance: considerations for captive breeding and reintroduction programs. Biological Conservation, 115(1), 71–77. 10.1016/S0006-3207(03)00095-8

Fischer, J., & Lindenmayer, D. B. (2000). An assessment of the published results of animal relocations. In Biological Conservation (Vol. 96, pp. 1–11).

Garnett, S. T., & Baker, G. B. (2021). The action plan for Australian birds 2020. CSIRO publishing.

Geering, D., & French, K. (1998). Breeding Biology of the Regent Honeyeater Xanthomyza phrygia in the Capertee Valley, New South Wales. Emu - Austral Ornithology, 98(2), 104–116. 10.1071/MU98011

Greggor, A. L., Berger-Tal, O., Blumstein, D. T., Angeloni, L., Bessa-Gomes, C., Blackwell, B. F., St Clair, C. C., Crooks, K., de Silva, S., & Fernández-Juricic, E. (2016). Research priorities from animal behaviour for maximising conservation progress. Trends in ecology & evolution, 31(12), 953–964.

Greggor, A. L., & Goldenberg, S. Z. (2023). Manipulating animal social interactions to enhance translocation impact. Trends in ecology & evolution, 38(4), 316–319.

Heinsohn, R., Lacy, R., Elphinstone, A., Ingwersen, D., Pitcher, B. J., Roderick, M., Schmelitschek, E., Van Sluys, M., Stojanovic, D., Tripovich, J., & Crates, R. (2022). Population viability in data deficient nomadic species: What it will take to save regent honeyeaters from extinction. Biological Conservation, 266. 10.1016/j.biocon.2021.109430

Hibbard, C. 2011, Regent Honeyeater Studbook, Australian Species Management Program, Zoo and Aquarium Association, Australia.

Hothorn, T., Bretz, F., Westfall, P., Heiberger, R. M., Schuetzenmeister, A., Scheibe, S., & Hothorn, M. T. (2016). Package ‘multcomp’. Simultaneous inference in general parametric models. Project for Statistical Computing, Vienna, Austria, 1–36.

IUCN. 2018. The IUCN Red List of Threatened Species. Version 2018-2. Available at: http://www.iucnredlist.org.

JMP, Version <17.2.0>, SAS Institute Inc., Cary, NC, 1989–2023

Kvistad, L., Ingwersen, D., Pavlova, A., Bull, J. K., & Sunnucks, P. (2015). Very low population structure in a highly mobile and wide-ranging endangered bird species. PLoS ONE, 10(12). 10.1371/journal.pone.0143746

Laiolo, P., Vögeli, M., Serrano, D., & Tella, J. L. (2008). Song diversity predicts the viability of fragmented bird populations. PLoS ONE, 3(3), e1822.

Lewis, R., Williams, L., & Gilman, R. (2020). The uses and implications of avian vocalizations for conservation planning. Conservation Biology, 35. 10.1111/cobi.13465

Lewis, R. N., Williams, L. J., & Gilman, R. T. (2021). The uses and implications of avian vocalizations for conservation planning. Conservation Biology, 35(1), 50–63. 10.1111/cobi.13465

Mackevicius, E. L., Gu, S., Denisenko, N. I., & Fee, M. S. (2023). Self-organization of songbird neural sequences during social isolation. Elife, 12, e77262.

Martins, B. A., Rodrigues, G. S. R., & de Araújo, C. B. (2018). Vocal dialects and their implications for bird reintroductions. Perspectives in ecology and conservation, 16(2), 83–89.

Mathews, F., Orros, M., McLaren, G., Gelling, M., & Foster, R. (2005). Keeping fit on the ark: assessing the suitability of captive-bred animals for release. Biological Conservation, 121(4), 569–577. 10.1016/j.biocon.2004.06.007

Mennill, D. J., Doucet, S. M., Newman, A. E., Williams, H., Moran, I. G., Thomas, I. P., Woodworth, B. K., & Norris, D. R. (2018). Wild birds learn songs from experimental vocal tutors. Current Biology, 28(20), 3273-3278. e3274.

Mets, D. G., & Brainard, M. S. (2018). Genetic variation interacts with experience to determine interindividual differences in learned song. Proceedings of the National Academy of Sciences, 115(2), 421–426.

Nottebohm, F., Nottebohm, M. E., & Crane, L. (1986). Developmental and seasonal changes in canary song and their relation to changes in the anatomy of song-control nuclei. Behavioral and neural biology, 46(3), 445–471.

Ogden, R., Chuven, J., Gilbert, T., Hosking, C., Gharbi, K., Craig, M., Al Dhaheri, S. S., & Senn, H. (2020). Benefits and pitfalls of captive conservation genetic management: Evaluating diversity in scimitar-horned oryx to support reintroduction planning. Biological Conservation, 241, 108244. 10.1016/j.biocon.2019.108244

Palagi, E., & Bergman, T. J. (2021). Bridging captive and wild studies: Behavioral plasticity and social complexity in Theropithecus gelada. Animals, 11(10), 3003.

Paxton, K. L., Sebastián-González, E., Hite, J. M., Crampton, L. H., Kuhn, D., & Hart, P. J. (2019). Loss of cultural song diversity and the convergence of songs in a declining Hawaiian forest bird community. Royal Society Open Science, 6(8). 10.1098/rsos.190719

Peters, S., Searcy, W. A., & Nowicki, S. (2014). Developmental stress, song-learning, and cognition. American Zoologist, 54(4), 555–567.

Price, T. D. (2002). Domesticated birds as a model for the genetics of speciation by sexual selection. Genetics of Mate Choice: From Sexual Selection to Sexual Isolation, 311–327.

R Core Team. (2020). R: A Language and Environment for Statistical Computing. In R Foundation for Statistical Computing https://www.R-project.org/

Robert, A. (2009). Captive breeding genetics and reintroduction success. Biological Conservation, 142(12), 2915–2922. 10.1016/j.biocon.2009.07.016

Ross, A. K., Letnic, M., Blumstein, D. T., & Moseby, K. E. (2019). Reversing the effects of evolutionary prey naiveté through controlled predator exposure. Journal of Applied Ecology, 56(7), 1761–1769.

Scheele, B., Hoffmann, E., & West, M. (2021). Recommendations for Conservation Translocations of Australian Frogs. NESP Threatened Sprecies Recovery Hub Project, 3(6).

Schloerke, B., Crowley, J., & Cook, D. (2018). Package ‘ggally’. Extension to ‘ggplot2, 713.

Searfoss, A. M., Pino, J. C., & Creanza, N. (2020). Chipper: Openℒsource software for semiℒautomated segmentation and analysis of birdsong and other natural sounds. Methods in Ecology and Evolution, 11(4), 524–531.

Seddon, P. J., Armstrong, D. P., & Maloney, R. F. (2007). Developing the Science of Reintroduction Biology. Conservation Biology, 21(2), 303–312. 10.1111/j.1523-1739.2006.00627.x

Seddon, P. J., Griffiths, C. J., Soorae, P. S., & Armstrong, D. P. (2014). Reversing defaunation: Restoring species in a changing world. Science, 345(6195), 406–412. 10.1111/1365-2664.12256

Shier, D. M., & Owings, D. H. (2007). Effects of social learning on predator training and postrelease survival in juvenile black-tailed prairie dogs, Cynomys ludovicianus. Animal Behaviour, 73(4), 567–577.

Shier, D. M., & Swaisgood, R. R. (2012). Fitness costs of neighborhood disruption in translocations of a solitary mammal. Conservation Biology, 26(1), 116–123.

Slabbekoorn, H., & Smith, T. B. (2002). Bird song, ecology and speciation. Philosophical Transactions of the Royal Society of London. Series B: Biological Sciences, 357(1420), 493–503.

Slade, B., Parrott, M. L., Paproth, A., Magrath, M. J. L., Gillespie, G. R., & Jessop, T. S. (2014). Assortative mating among animals of captive and wild origin following experimental conservation releases. Biology Letters, 10(11), 20140656. 10.1098/rsbl.2014.0656

Soma, M. F. (2011). Social factors in song learning: a review of Estrildid finch research. Ornithological Science, 10(2), 89–100.

Taylor, G., Ewen, J. G., Clarke, R. H., Blackburn, T. M., Johnson, G., & Ingwersen, D. (2018). Video monitoring reveals novel threat to critically endangered captive-bred and released regent honeyeaters [Article]. Emu, 118(3), 304–310. 10.1080/01584197.2018.1442227

Teitelbaum, C. S., Converse, S. J., & Mueller, T. (2019). The importance of early life experience and animal cultures in reintroductions. Conservation Letters, 12(1), e12599. 10.1111/conl.12599

Thorpe, W. H. (1958). The learning of song patterns by birds, with especial reference to the song of the chaffinch Fringilla coelebs. Ibis, 100(4), 535–570.

Tripovich, J. S., Popovic, G., Elphinstone, A., Ingwersen, D., Johnson, G., Schmelitschek, E., Wilkin, D., Taylor, G., & Pitcher, B. J. (2021). Born to Be Wild: Evaluating the Zoo-Based Regent Honeyeater Breed for Release Program to Optimise Individual Success and Conservation Outcomes in the Wild. Frontiers in Conservation Science, 2, 16–16. 10.3389/fcosc.2021.669563

Varkevisser, J. M., Mendoza, E., Simon, R., Manet, M., Halfwerk, W., Scharff, C., & Riebel, K. (2022). Multimodality during live tutoring is relevant for vocal learning in zebra finches. Animal Behaviour, 187, 263–280.

Vecsei, M. (2015). Juvenile Song Learning in Regent Honeyeaters, Anthochaera phrygia, at Taronga Zoo, Australia. Sydney, NSW: Faculty of Science, Department of Biological Sciences Macquarie University, Sydney, NSW Australia in partial fulfilment of the requirements for the degree of Master of Research.

Viggers, K., Lindenmayer, D., & Spratt, D. (1993). The Importance of Disease in Reintroduction Programmes. Wildlife Research, 20(5), 687–698. 10.1071/WR9930687

Whiten, A. (2021). The burgeoning reach of animal culture. In Science (Vol. 372): American Association for the Advancement of Science.

Williams, J., & Slater, P. (1990). Modelling bird song dialects: the influence of repertoire size and numbers of neighbours. Journal of Theoretical Biology, 145(4), 487–496.

Williams, S. E., & Hoffman, E. A. (2009). Minimizing genetic adaptation in captive breeding programs: A review. Biological Conservation, 142(11), 2388–2400. 10.1016/j.biocon.2009.05.034

