## Supplementary Information for "Reintroduction of wild song culture to a critically endangered songbird"

**Supplementary Materials**

**Supplementary Text**

**Text 1 – Rationale for adaptive decisions for song tutoring protocols.**

In the 2020/21 season (year one), we trained one cohort with playback at Taronga Western Plains Zoo (TWPZ), one cohort with live tutors at Taronga Zoo (TZ) and one cohort remained un-tutored as a control group at TWPZ. It was clear that live tutoring resulted in better song outcomes than playback, while playback alone was only marginally better the control group. Songs from the live tutoring only (Large cohort) did appear to be more complex than the playback and control groups, however, the final songs of all groups did not approach the wild reference song from a listener’s perspective and subsequent statistical analysis of acoustic features confirmed this (as reported in the main text). This evidence, combined with observations from researchers and husbandry staff showed that these methods had potential if some minor changes were made to the protocols.

During the first season, we observed that in the playback group, the vocalisations of the juveniles we far more frequent than the playback itself, and that juvenile birds tended to interact and vocalise more often in response to each other than in response to playback emitted from the loudspeakers. We believed that interactions with real birds are more enriching than playback and as such juveniles would preferentially learn from a live model, even if they were also juvenile (i.e., Varkevisser 2022; Derregnaucourt 2013). A similar pattern of behaviour was observed in the live tutor group, whereby juveniles did interact with the tutor, but the volume of juveniles appeared to result in most interactions being with other juveniles instead of the adult tutor. We also noted that for the earliest period of their life, young birds didn’t hear wild song as the tutor was not vocal during his own breeding period. This resulted in the following changes for season 2021/22:

(1) Rate of song playback was doubled in the both the playback only and live tutoring cohorts.
(2) Adult tutor to juvenile ratio was limited to a maximum of1:5.
(3) Playback was introduced during the early phases of the live tutoring cohort. The previous live tutoring cohort in 2021 was not exposed to additional playback.

Given the severity of ongoing population decline in regent honeyeaters and the importance of the zoo-breeding program to their recovery efforts (Heinsohn *et al.,* 2022; Appleby *et al.,* 2024), we prioritised implementing all the above changes at once to maximise the chances of successful song learning over a more rigorous step-by-step experimental approach. This was also in-part driven by a lack of space in the zoo setting, where implementing each experimental change in separate cohorts would have taken several years to deploy.

In 2021/22 the playback only (small) cohort showed little improvement compared with the playback only (large) cohort. This indicated that song learning is likely multimodal (Varkevisser et al. 20220) in regent honeyeaters; that is, interaction with a tutor is critical and as such, we decided to cease playback only cohorts in future years in favour of expanding live tutoring. In the live tutoring + playback cohort, we frequently detected wild-type songs by ear during the song tutoring which was later confirmed statistically and are displayed below in fig. S1.

While the 2021/22 season produced juvenile birds whose songs resembled the reference wild song type, the decision to implement multiple changes to the protocols in one year created ambiguity around whether the smaller cohort or the reinforcement of the tutor with playback was the critical factor determining successful song learning. To disambiguate this, we introduced the live tutoring only (small) cohort.

**Supplementary Text 2 – Composing the Playback Track**

We first gathered 25 recordings of wild-origin males who sang the Typical Blue Mountains song-type. These were sourced largely from the wild using recordings obtained in Crates et. al (2021, n = 23), as well as one recording made by D. A of a wild-origin bird at TZ who sang Blue Mountains typical song-type and one recording from Xeno-Canto.

We initially composed 14 playback tracks. Each day of the week had a unique playback track, and two time periods had a set of seven tracks. The first seven of these time periods were 12 hours in length to fill the average Austral daylight hours between September and November. The second seven were 14 hours in duration, designed to fill the average Austral daylight hours between December and March.

We divided tracks between peak and off-peak periods. The peak period was designed with the intention of mimicking the increased call activity of songbirds early in the morning and late evening periods (Catchpole and Slater, 2008). During the peak period, songs were broadcast at a more frequent rate than the off-peak period during the middle of the day. On our tracks the peak period ran for the first four, and the last two hours of the day, regardless of day length. The off-peak period ran in the warmer, middle part of the day in between peak periods. The off-peak period of the playback track broadcast songs at a less frequent rate than the peak period. Between September and November, the off-peak period was six hours in length. Between December and March, the off-peak period was eight hours in length.

To compose the tracks, we developed two randomisation matrices each for the peak and off-peak periods. The first randomisation matrix was designed to set the order of the song presentation and then the timing of the song presentations. For the peak period, songs were emitted at an average rate of one song per 25 seconds (randomised silence between 12- 75 seconds). For the off-peak period, songs were emitted at an average rate of one song per 93 seconds (randomised silence between 60-180 seconds).

We first post-processed recordings in audacity to remove any unwanted sounds. We then ran a 300Hz High Pass Filter to all songs to reduce low end noise. We then created 10, one-hour long blocks of both peak and off-peak playback, by arranging songs in Apple’s ‘garageband’ according to the randomisation matrix. We created another randomisation matrix to decide the presentation of one-hour blocks to make the peak and off peak tracks for each day. These one-hour blocks were arranged in apple’s ‘garageband’ according to the randomisation matrix to make seven peak periods of four hours (morning) and seven peak periods of two hours (evening). We repeated this to make seven off peak periods of six hours of length, and seven off peak periods of eight hours in length. To compose the final 12- and 14-hour playback tracks we randomly assigned one morning peak period, one off peak period and one evening off peak period to create one track. This was done in Audacity software owing to the extremely large file size that was inappropriate for garage band. Files were exported as high-quality ‘.wav’ files and assigned one day each per week for broadcast.

In 2021/2022 season we increased the song playback rate. To do this we followed all of the above steps this time decreasing the randomised silence period by half (effectively doubling the playback rate).

**Supplementary Text 3 – Playback Systems**

We employed two playback systems throughout the three years. At both zoos the broadcast speakers were Bose ‘Freespace’ outdoor speakers. In TZ we used a XX amplifier connected to a windows computer. Playback was initiated using the ‘Windows Task Scheduler’ software. At TWPZ, where a permanent indoor space was not available, we used a portable set up housed in a waterproof box. We used a Swamp Industries line amplifier connected to an iPod touch. We used the default automation app to schedule playback.

**Supplementary Tables**

***Supplementary Table 1*** –

Timeline Example 2022/23 season (Live Tutor + Playback). Table shows the timeline for two live tutoring + playback cohorts in the 2022/23 season. Light-blue shading represents the period where the juvenile bird is in the natal aviary with its parents. Pale-yellow represents the period when the bird is in a an experimental creche exposed to playback while next to a wild male with who they can’t physically interact but can see and hear. The pale-green shading represents the period when birds are in the creche aviary with the live tutor as well as exposed to song playback. The dashed red line denotes the addition of the live to tutor to the creche aviary.

| ID | August | | | September | | | | October | | | | November | | | | December | | | | January | | | | February | | | | March | | | |
| --- | --- | --- | --- | --- | --- | --- | --- | --- | --- | --- | --- | --- | --- | --- | --- | --- | --- | --- | --- | --- | --- | --- | --- | --- | --- | --- | --- | --- | --- | --- | --- |
| **WEEK** | 3 | 4 | | 1 | 2 | 3 | 4 | 1 | 2 | 3 | 4 | 1 | 2 | 3 | 4 | 1 | 2 | 3 | 4 | 1 | 2 | 3 | 4 | 1 | 2 | 3 | 4 | 1 | 2 | 3 | 4 |
| MALE | BIRD 1 BORN | |  |  |  |  |  |  |  |  |  |  |  |  |  |  |  |  |  |  |  |  |  |  |  |  |  |  |  |  |  |
| MALE |  | |  |  | BIRD 3 BORN | |  |  |  |  |  |  |  |  |  |  |  |  |  |  |  |  |  |  |  |  |  |  |  |  |  |
| MALE |  | |  |  |  |  |  |  |  | BIRD 4 BORN | |  |  |  |  |  |  |  |  |  |  |  |  |  |  |  |  |  |  |  |  |
| MALE |  | |  |  |  |  |  |  |  | BIRD 5 BORN | |  |  |  |  |  |  |  |  |  |  |  |  |  |  |  |  |  |  |  |  |
| MALE |  | |  |  |  |  |  |  |  |  |  |  |  | BIRD 8 BORN | |  |  |  |  |  |  |  |  |  |  |  |  |  |  |  |  |
| MALE |  | | BIRD 2 BORN | |  |  |  |  |  |  |  |  |  |  |  |  |  |  |  |  |  |  |  |  |  |  |  |  |  |  |  |
| MALE |  | |  |  |  |  |  | BIRD 3 BORN | |  |  |  |  |  |  |  |  |  |  |  |  |  |  |  |  |  |  |  |  |  |  |
| MALE |  | |  |  |  |  |  |  |  |  | BIRD 6 BORN | |  |  |  |  |  |  |  |  |  |  |  |  |  |  |  |  |  |  |  |
| MALE |  | |  |  |  |  |  |  |  |  |  | BIRD 7 BORN | |  |  |  |  |  |  |  |  |  |  |  |  |  |  |  |  |  |  |

**Supplementary Table 3:** Acoustic features measured in ‘Chipper’ and whether they were included in the discriminant function analysis.

| **Feature** | **Included in DFA?** |
| --- | --- |
| Number of Unique Syllables | **Yes** |
| Smallest Syllable Frequency Range (Hz) | **Yes** |
| Average Syllable Lower Frequency (Hz) | **Yes** |
| Largest Syllable Frequency Range (Hz) | **Yes** |
| Number Syllables per Bout Duration (1/ms) | **Yes** |
| Number of Syllables | **Yes** |
| Overall Syllable Frequency Range (Hz) | **Yes** |
| Std. Deviation Syllable Frequency Range (Hz) | **Yes** |
| Smallest Syllable Duration (ms) | **Yes** |
| Average Silence Duration (ms) | **No** |
| Average Syllable Duration (ms) | **No** |
| Average Syllable Duration (ms) | **No** |
| Average Syllables Upper Frequency (Hz) | **No** |
| Bout Duration(ms) | **No** |
| Largest Silence Duration (ms) | **No** |
| Largest Syllable Duration (ms) | **No** |
| Largest Syllable Freq Range (Hz) | **No** |
| Max Syllables Frequency (Hz) | **No** |
| Min Syllables Frequency (Hz) | **No** |
| Smallest Silence Duration (ms) | **No** |
| Std. Dev. Silence Duration (ms) | **No** |

**Supplementary Table 3**

| Table showing results of backwards model selection. Each step removes a predictor and shows the resulting AIC. Text in bold shows the strongest performing model. | | | |
| --- | --- | --- | --- |
| Model Formula | AIC | Predictor to Remove | New AIC |
| Mahalanobis Distance~ Treatment + Age | 264.71 | Age | 262.74 |
|  |  | Treatment | 334.25 |
| **Mahalanobis + Distance~ Treatment** | **262.7** | Treatment | 340.3 |

**Supplementary Table 4**

Table showing the model estimates of generalised linear model regressions using to formula Mahalanobis distance ~ Treatment + Age

|  | Mahalanobis Distance | | |
| --- | --- | --- | --- |
| *Predictors* | *Estimates* | *CI* | *p* |
| (Intercept) | 2.56 | 0.13 – 4.99 | **0.039** |
| Control (T1) | 2.03 | 0.40 – 3.65 | **0.014** |
| Playback (Large, T2) | 1.22 | -0.32 – 2.77 | 0.121 |
| Live Tutoring Only (Large, T3) | 2.55 | 1.16 – 3.94 | **<0.001** |
| Playback (Small, T4) | 2.30 | 0.90 – 3.70 | **0.001** |
| Live Tutoring + Playback (T5) | -0.33 | -1.74 – 1.08 | 0.643 |
| Live Tutor (Small, T6) | -0.22 | -1.75 – 1.30 | 0.773 |
| Age | -0.00 | -0.01 – 0.01 | 0.854 |

**Supplementary Figures**

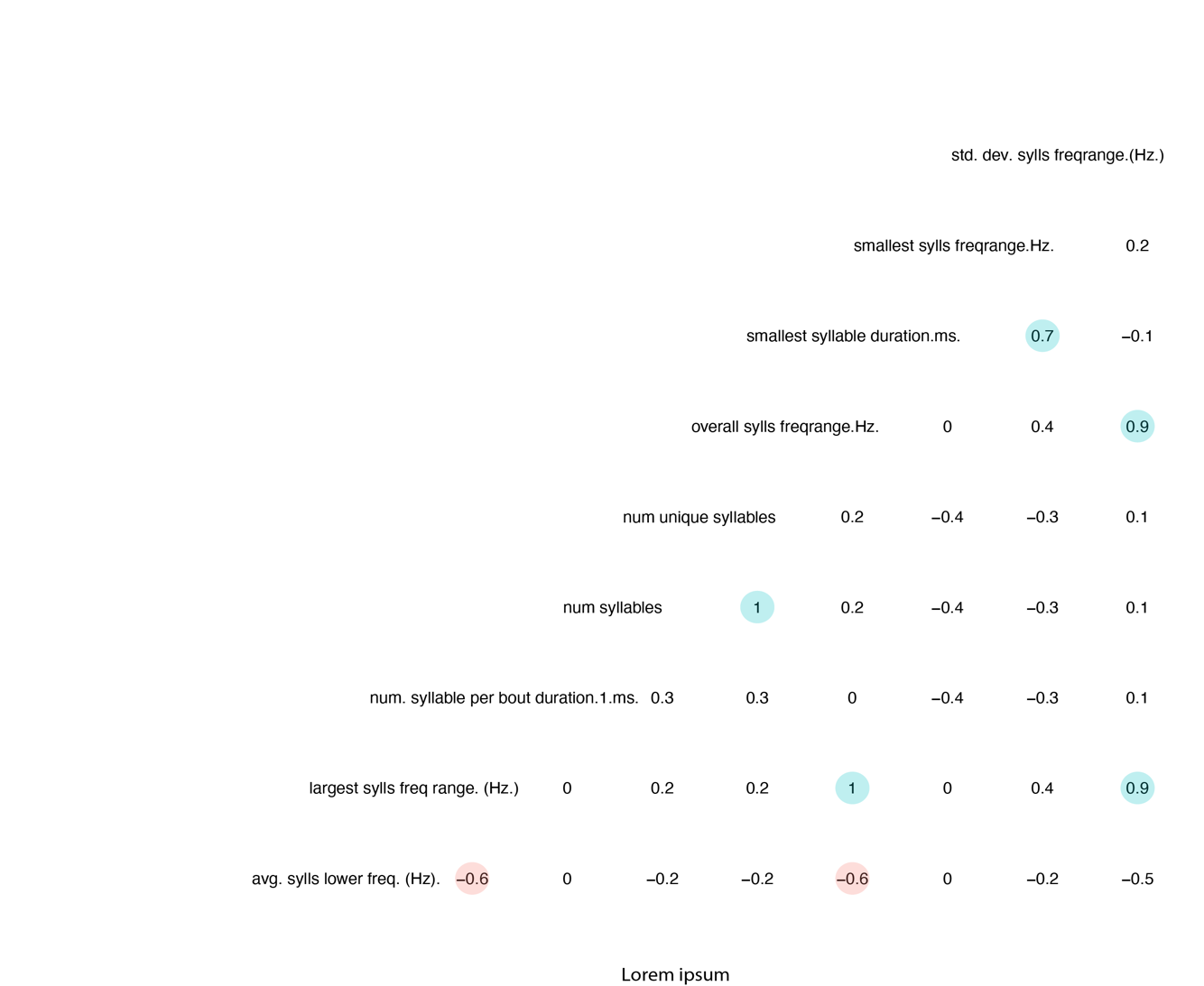

Fig. S1 – Correlation matrix of acoustic features included in discriminant function analysis.

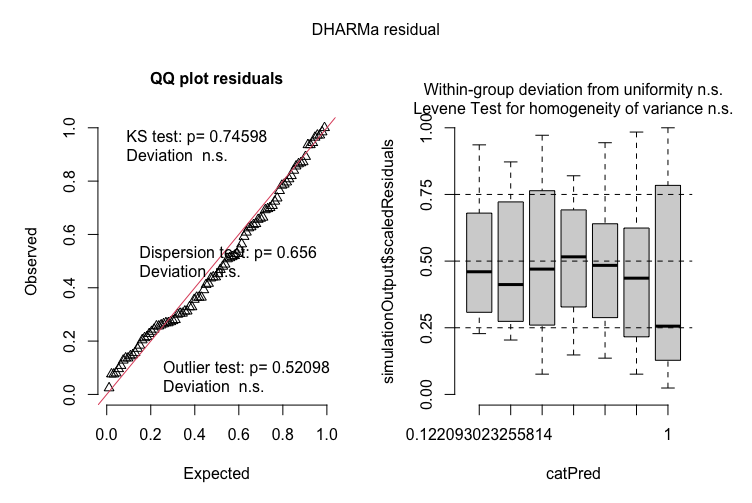

Fig S2. Simulated Residuals of Generalised Linear Regression (Mahalanobis Distance ~ Treatment)
